# Hierarchical prediction and perturbation of chromatin organization reveal how loop domains mediate higher-order architectures

**DOI:** 10.1101/2025.05.25.656045

**Authors:** Jiachen Wei, Yue Xue, Yi Qin Gao

## Abstract

The genome is folded within the dense cell nucleus in a hierarchical manner, resulting in complex interactions between distinct folding strategies at various length scales. To elucidate how short-range loop domains regulate higher-order structures of the chromatin, such as topologically associating domains (TADs) and compartments, we introduce HiCGen, a hierarchical and cell-type-specific generator based on Swin-transformer architecture. HiCGen predicts genome organization across different spatial scales utilizing DNA sequence and genomic features as inputs. The model enables *in silico* screening through genetic or epigenetic perturbations on genome architecture, with resolution down to 1 kb. Our analysis reveals unexpected linear correlations between loop properties and genome organization at various levels, including insulation degree, compartmentalization, and contact intensity over genomic distances exceeding 10 Mb. Regional or global perturbation conducted by HiCGen provides biological implications for such cross-scale correlations and their genome-function dependence. Notably, perturbation analysis of the human genome in sigmoid colon tissue demonstrates that modest activation of carcinogenesis-associated enhancers is sufficient to hijack nearby promoter, reshape TAD boundaries, and even flip compartment at mega-base scale.

## Introduction

The genome is organized within the cell nucleus in a complex, hierarchical manner, which is crucial for regulating gene expression and maintaining cellular state and functions^1–6^. This hierarchical organization spans from the chromatin loops at kilobase scale to topologically associating domains (TADs), and compartments at megabase scale. Although loops are much smaller in size compared to higher-order chromosome architectures, recent studies^7–15^ have noticed their correlations with insulation degree, compartmentalization, as well as long-range contact intensity. Understanding the interplay between different hierarchies is essential for elucidating the mechanisms of genome function and regulation.

The high-throughput chromosome conformation capture (Hi-C) contact maps have revolutionized the understanding of genome organization by providing comprehensive information of chromatin interactions ^3,16^. Hi-C captures the frequency of contacts between different genomic loci, revealing the spatial organization of the genome at various scales. Through Hi-C, one can identify loops, TADs and compartments, thereby uncovering the hierarchical nature of genome organization.

Many computational techniques have been developed to interpret cell-type-specific Hi-C matrices. For instance, polymer models that had quantitative alignment to Hi-C and other biological data ^17–19^ were designed to understand genomic organization. Deep learning has emerged as a powerful technique for integrating multi-modal data, capturing intricate, non-linear interactions among DNA sequence, epigenetic marks, and genome structure ^20^. Several deep neural networks have been developed recently to predict Hi-C contact maps based solely on genome sequence or together with other genomic features. For instance, Akita ^21^

and DeepC ^22^ have a similar convolutional module, both of which take DNA sequence as input and can only predict interactions within a 1-Mb window. Orca ^23^ is the first architecture that captures the hierarchical organizations of the genome from 4-kb up to 1-Mb resolution. However, as a sequence-based convolutional network, Orca can only predict contact maps for the trained cell line and may suffer from pattern loss. Taken both sequence and genomic features as inputs, C.Origami ^24^ enables *de novo* predictions of the genome organization across cell types. The model is based on a encoder-decoder architecture, enabling a 2-Mb window of prediction at 8-kb resolution. However, because of the fixed output resolution, C.Origami can barely characterize the complex interplay between different folding strategies at distinct length scales.

To address these limitations, we introduce HiCGen, a hierarchical and cell-type-specific generator based on Swin-transformer architecture^25,26^. HiCGen predicts cell-type-specific genome organizations across spatial scales (i.e. from 1-kb to 128-kb resolutions) by integrating DNA sequence and genomic features as inputs. The model outperforms previous methods for predicting genomic contacts, in terms of accuracy, multiscale structure (i.e. loops, TADs and compartments) characterization, and generalization across cell types. The model also facilitates *de novo* prediction of gene-edited samples, and high-throughput screening through perturbations of genetic or epigenetic signals. By leveraging the power of deep learning, the model provides new insights into the interplay of hierarchical genome organization, and, in particular, specific interactions between enhancers and promoters^27–30^.

## Results

### Predictions of the hierarchical genome interactions

**Fig. 1a** illustrates the schematic architecture of HiCGen. The input features include one-hot encoding of the genomic sequence, along with ATAC-seq and CTCF ChIP-seq profiles. In addition to the convolutional neural network (CNN) used for encoding and decoding, HiCGen leverages the Swin-transformer architecture with up- sampling techniques (Methods) to capture the hierarchy of contact matrices^25,26^. By integrating information from contact maps of broader interaction ranges, the model is able to predict genomic structures across 128-fold resolution range.

**Fig. 1.**
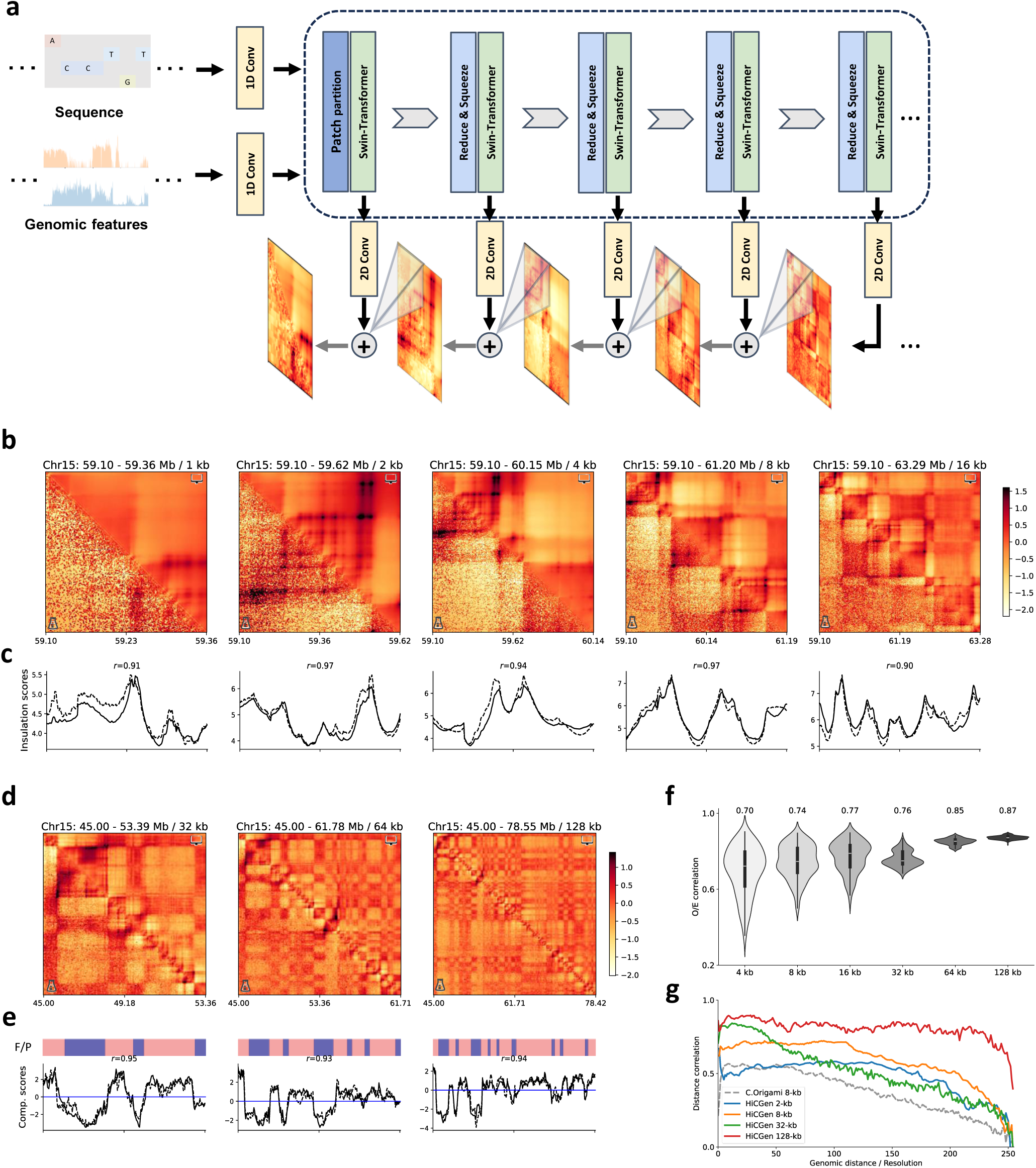
| Predictions of hierarchical genome organization by HiCGen. a, Schematic diagram illustrating the architecture of HiCGen. **b,c,** Comparison of experimental (lower triangular) and predicted (upper triangular) contact maps for a genomic region from the test set (chr15) of GM12878 at 1-kb, 2-kb, 4-kb, 8-kb and 16-kb resolutions (**b**), along with experimental (dashed lines) and predicted (solid lines) insulation scores (**c**) for the corresponding maps. **d,e,** Comparison of experimental and predicted contact maps for a region randomly selected from test set at 32-kb, 64-kb and 128-kb resolutions (**d**), accompanied by the CGI forest/prairie domains (F/P), experimental (dashed) and predicted (solid) compartment scores (**e**) for the corresponding maps. **f** Comparisons of Pearson correlation coefficients between experimental and predicted contacts at 4-to-128-kb resolutions for chr15. **h,** Distance-stratified correlations between experimental and predicted matrices of chr15 across various resolutions. The dashed line indicates predictions of C.Origami of its test set at 8-kb resolution. Experimental contact maps are represented in log fold over distance-based background without adaptive coarse-graining.

**Fig. 1b** compares the hierarchical contact maps derived from intact Hi-C experiment of GM12878 with those predicted by the model at 1-16-kb resolutions. Certain loops, possibly due to their brief formation periods ^31^, are challenging to capture via Hi-C. While most loops, stripes, and TAD boundaries are accurately predicted by HiCGen, the model is also capable of identifying loops that are faint or absent in experimental data at high resolution (see also Supplementary Fig. 8). HiCGen also effectively captures protein-mediated interactions and promoter-enhancer interactions (Supplementary Fig. 8). The insulation scores calculated from experimental Hi-C exhibit strong agreement with that derived from predicted matrices, with a Pearson correlation *r*≃0.95 (**Fig. 1c**). **Fig. 1d** compares experimental and predicted contact maps at 32-128-kb resolutions. HiCGen accurately captures compartment-level structures: the compartment scores derived from experimental Hi-C also align well with those from predicted contact matrices, with Pearson correlation *r*≃0.94 (**Fig. 1e**).

**Figs. 1f** presents correlations between experimental and predicted contact maps across different resolutions for the test-set chromosome (i.e. chr15). Due to effective encoding of multi-scale information, relatively high correlations (*r*>0.7) are achieved by HiCGen. **Fig. 1g** compares the distance-stratified correlations between experimental and predicted data for chr15. HiCGen demonstrates higher distance-stratified correlations at each resolution compared to C.Origami at 8-kb resolution. The results demonstrate the robust capability of HiCGen in predicting multiscale genome structures.

### Cross-cell-type predictions

By incorporating cell-type-specific CTCF ChIP-seq and ATAC-seq signals as inputs, HiCGen enables *de novo* predictions of chromatin organization for previously unseen cell types. **Fig. 2a-b** compares experimental and predicted contact maps for a new cell type, IMR90, across different resolutions. Despite being trained exclusively on GM12878, the model accurately recapitulates genome-wide interaction patterns, including IMR90-specific ones. Regions with CTCF ChIP-seq peaks or abrupt changes in ATAC-seq profiles (indicated by black arrows) — hallmarks of loop anchors and TAD boundaries — are faithfully reproduced in the predictions. The predicted TAD boundaries and compartments align well with those derived from experimental data, as evidenced by Pearson correlation coefficients *r*≃ 0.9 for insulation and *r*≃ 0.8 for compartment scores. We further applied the model to predict contact maps of HCT116, a colon cancer cell line (**Fig. 2c-d**). While interactions at compartment level are comprehensively captured, the model seems to underestimate the short-range interactions at TAD level in HCT116. In fact, contacts at loop/TAD levels are enhanced for most cancer cells compared to that for normal tissues 30,31. These short-range interactions can be well captured by model trained with other type of cancer cell lines (Supplementary Fig. 9).

**Fig. 2.**
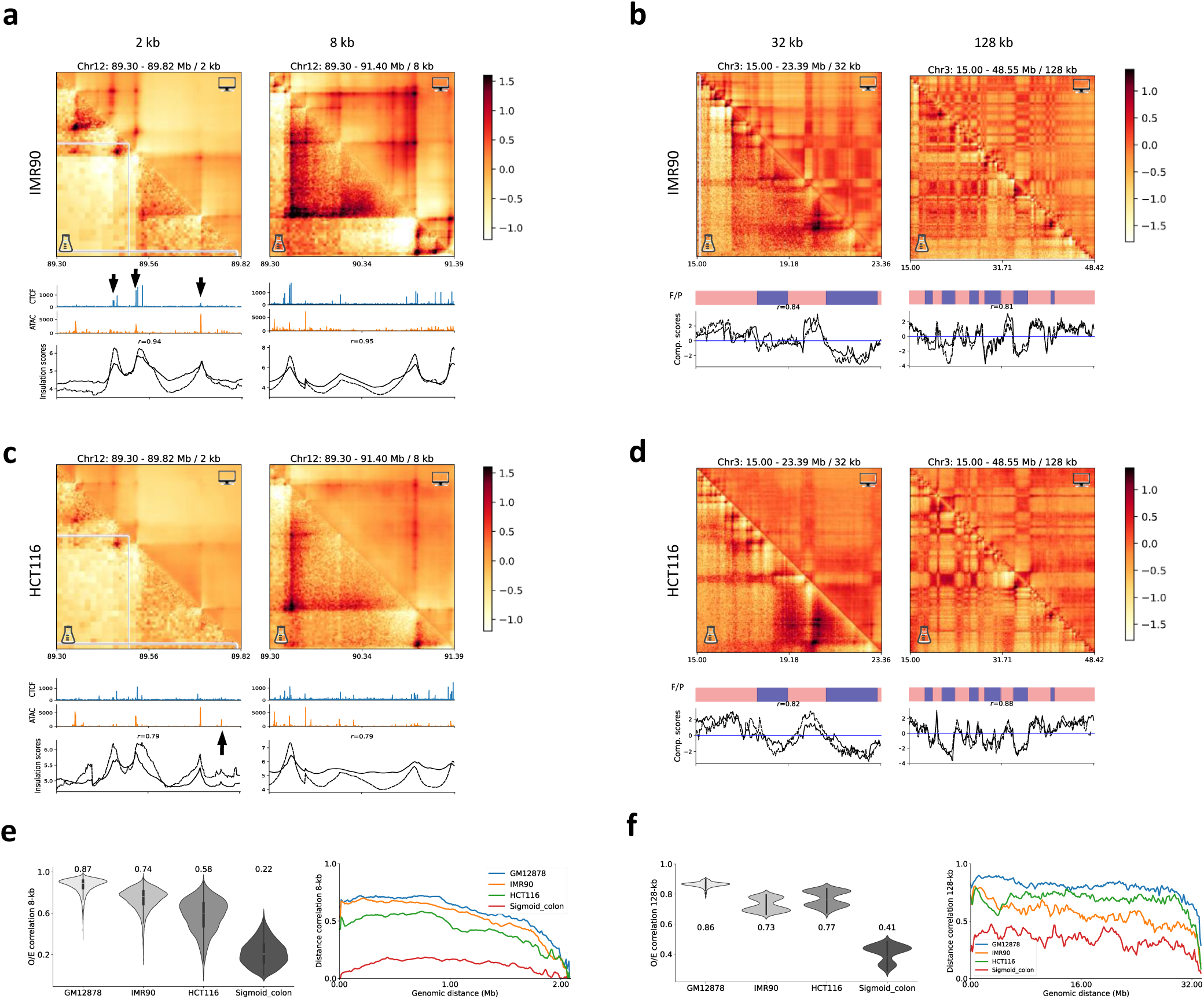
| Cross-cell-type predictions of genomic contacts. a,c, Comparison of experimental (lower triangular) and predicted (upper triangular) contact maps for a genomic region of IMR90 (**a**) and HCT116 (**c**) at 2-kb and 8-kb resolutions, along with genomic features as inputs, experimental (dashed lines) and predicted (solid lines) insulation scores for the corresponding maps. **b,d,** Comparison of experimental and predicted contact maps for a selected genomic region of IMR90 (**b**) and HCT116 (**d**) at 32-kb and 128-kb resolutions, along with corresponding CGI forest/prairie domains (F/P), compartment scores derived from experimental data (dashed lines) and predictions (solid lines). **e,f,** Comparisons of Pearson correlations (left panel) and distance-stratified correlations (right panel) between experimental and predicted contacts at 8-kb (**e**) and 128-kb (**f**) resolutions. Contact maps are represented in log fold over distance-based background. All metrics and contact maps are derived from the predictions of a model trained with GM12878 cell line.

The performance of the model trained with GM12878 in predicting different types of cells are presented in **Fig. 2e-f**. Compared to that for IMR90 and HCT116, poor correlations between predicted and experimental data were observed for sigmoid colon. Nevertheless, we find the model performance (on colon) can be significantly improved by training with other type of human tissues (Supplementary Fig. 9). The results demonstrate the ability of HiCGen in cross-cell-type prediction through rational selection of training and targeted samples.

### Predictions of genome reorganization upon genetic modifications

To evaluate the model performance on gene-edited samples, we conducted cross-cell-type predictions of unseen contact maps of HCT116 and its genetically modified counterparts HCT116^ΔMED14^, HCT116^ΔCDK7^ and HCT116^ΔCTCF^, in which MED14, CDK7 and CTCF were respectively abolished through auxin inducible degron^33^. **Fig. 3a-c** respectively present Hi-C contact maps and predicted matrices by model trained on PANC- 1 cell line at 2-kb, 8-kb and 32-kb resolutions. Marked alterations in CTCF ChIP-seq and ATAC-seq profiles (as shown in Fig. 3a), indicate substantial genome reorganization (e.g., dysfunctional extrusion boundaries in HCT116^ΔCTCF^). Despite these perturbations, the predicted matrices show good consistency with experimental maps for both HCT116 and its variants. The insulation and compartment scores obtained from Hi-C data of gene-edited samples also agree well with those derived from model predictions, yielding Pearson correlation coefficients *r*≃ 0.8 across different resolutions.

**Fig. 3.**
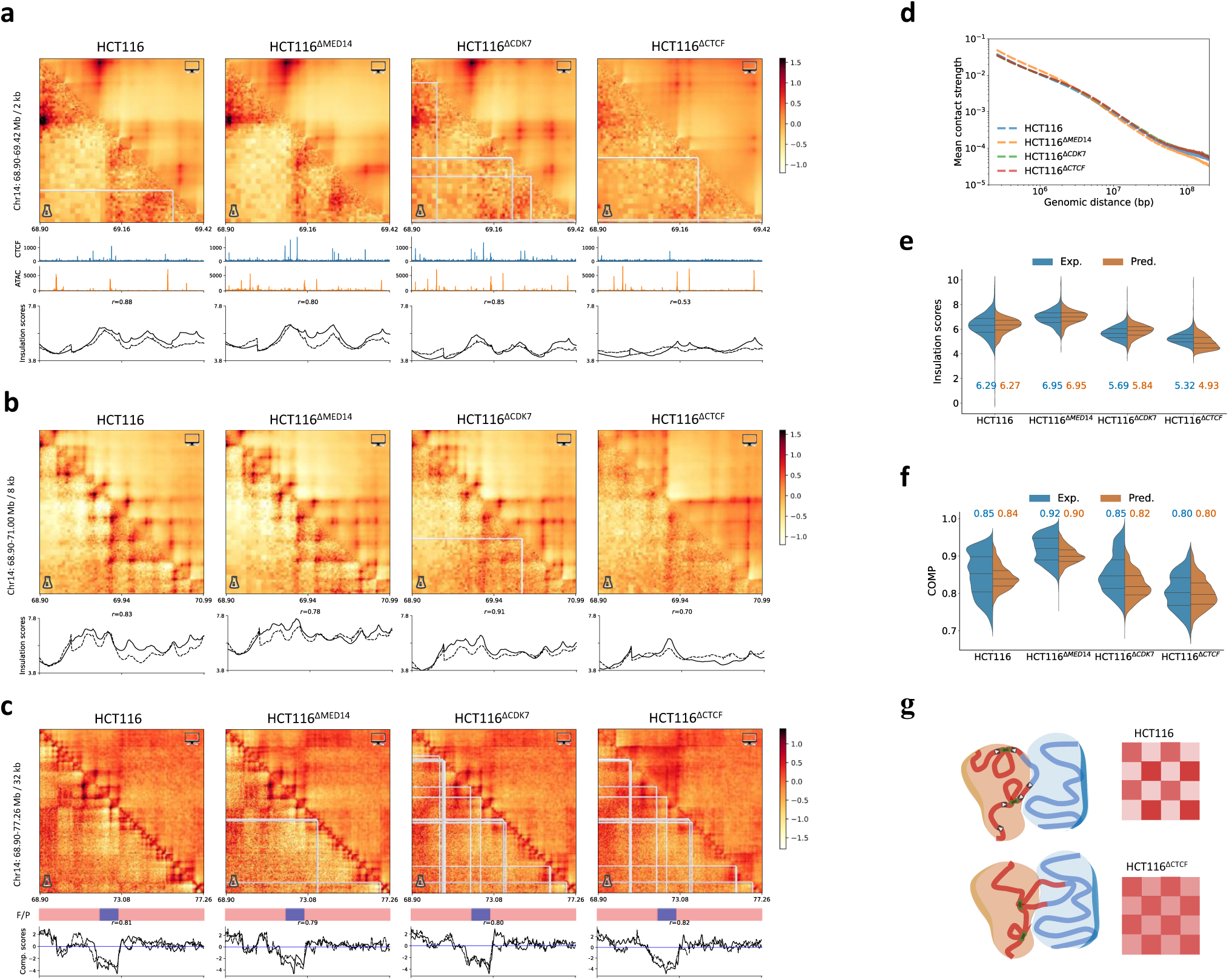
| Cross-scale predictions of gene-edited samples. a-c, Comparison of experimental (lower triangular) and predicted (upper triangular) contact maps for a selected genomic region of HCT116 and its gene-edited variants, HCT116^ΔMED14^, HCT116^ΔCDK7^ and HCT116^ΔCTCF^, at 2-kb (**a**), 8-kb (**b**) and 32-kb (**c**) resolutions, along with genomic features as inputs (**a**), experimental (dashed lines) and predicted (solid lines) insulation scores (**a,b**), CGI forest/prairie domains (F/P) and compartment scores (**c**) for the corresponding maps. **d,** Contact probability as a function of genomic distance for HCT116 and its gene-edited variants. **e,f,** Comparison of experimental and predicted insulation scores on all TAD boundaries at 8-kb resolution (**e**) and genome-wide compartment scores at 128-kb resolution (**f**) for HCT116 and its gene-edited variants. All metrics and contact maps are derived from the predictions of a model trained on PANC-1. **g,** Schematic diagram of how the loss of loop extrusion boundaries disrupts compartmentalization for HCT116.

**Fig. 3e** compares the experimental and predicted insulation scores at TAD boundaries for HCT116 and its genetically modified counterparts, indicating the slight increase of insulation degree upon MED14 knockout and the reduction of insulation degree upon CDK7 or CTCF knockout. Slight increase in compartmentalization degree (COMP) is observed in HCT116^ΔMED14^ relative to HCT116, while decreased COMP are observed for HCT116^ΔCDK7^ and HCT116^ΔCTCF^, as depicted in **Fig. 3f**. Both TAD and compartment-level fluctuations can be well captured by HiCGen, indicating the model perceives the cross-scale correlations regarding genetic modification. The decrease of COMP in HCT116^ΔCTCF^ may be attributed to the disruption of extrusion boundaries, as illustrated in **Fig. 3g**. This disruption can facilitate the formation of unregulated loops, resulting in increased contacts across adjacent compartments.

### *In silico* perturbation reveals cross-scale correlations

HiCGen enables *in silico* perturbation on regulatory elements through activation/deactivation of corresponding input signals. As illustrated in **Fig. 4a**, we screened the 1-kb fragments of the HCT116 genome and determined difference in contact maps across various scales upon silencing the corresponding ATAC-seq signals. The effect of silencing was quantified by the impact score, calculated as the mean absolute of the difference maps. To investigate the effect of perturbation on genome organization across multiple genomic scales, we further categorized the screened 1-kb fragments into gene bodies, promoters, enhancers, and other regions (Methods). **Fig. 4b** compares the composition of top 10% fragments with highest impact scores (HIS- subset) following the silencing of 1-kb ATAC-seq signals for HCT116 and HCT116^ΔCTCF^. The HIS-subset for HCT116^ΔCTCF^ exhibits greater enrichment of promoter regions than that for HCT116, suggesting potential activation of silenced promoters following CTCF knockout. These HIS-subset promoters in HCT116^ΔCTCF^ may experience stronger enhancer-promoter interactions than that in HCT116, as evidenced by higher relative enrichment of H3K27ac marks (Supplementary Fig. 15). Interestingly, the composition of regulatory elements within HIS-subset is barely changed across all analyzed scales (256 kb to 32 Mb; **Fig. 4b**), suggesting the multiscale regulatory influence of these elements on chromatin architecture.

**Fig. 4.**
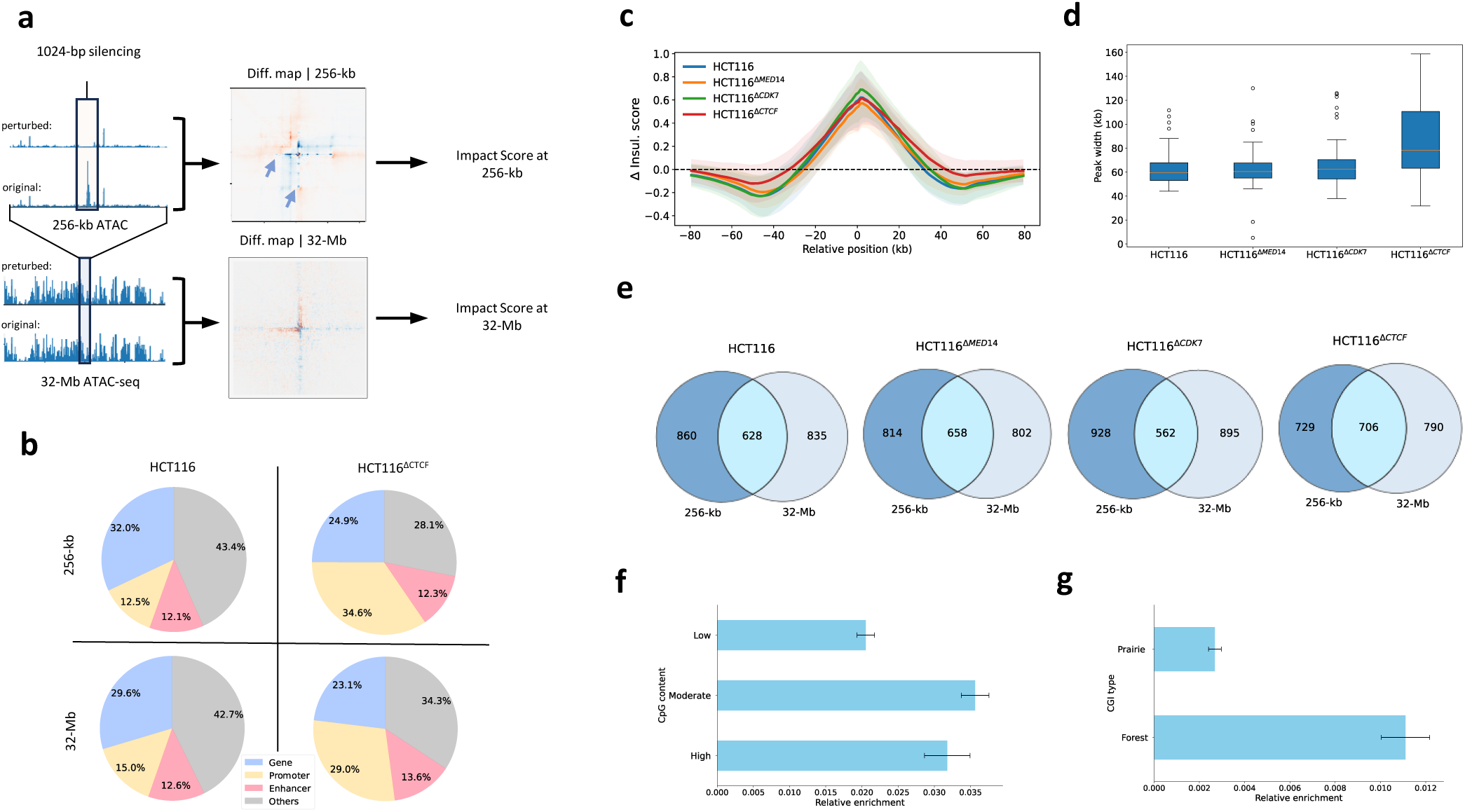
| *In silico* perturbation analysis of HCT116 through silencing active genome fragments. a, Workflow of screening out genome fragments with high impact to genome organizations: the genome was delimited into 1-kb fragments, and differences in contact matrices upon silencing the corresponding 1-kb ATAC-seq inputs were determined at each resolution with a sliding window centering at the perturbed region. Impact score was quantified as mean value of the absolute difference at each scale. **b,** Compositions of regulatory elements of the top 10% fragments with highest impact scores (HIS-subset) at smallest (256-kb) and largest (32-Mb) scales for HCT116 and HCT116^ΔCTCF^. **c,** The average differential insulation curves for HIS-subset at 1-kb resolution following the silencing of 1-kb ATAC- seq signals. Only portions of regulatory elements that experience enhanced insulation scores (i.e. with positive peaks) subsequent to the silencing were displayed. **d,** boxplot distributions of peak width of the differential insulation curves at 1-kb resolution for HIS-subset of HCT116 and its gene-edited variants, HCT116^ΔMED14^, HCT116^ΔCDK7^ and HCT116^ΔCTCF^. **e,** Venn diagrams of enhancers within HIS-subset at 256-kb and 32-Mb scales. **f,g,** Comparisons of CpG content levels (**f**) at transcription start site and CGI types (**g**) of promoters corresponding to overlapped enhancers with cross-scale impact obtained in (**e**). The results suggest pivotal regulatory elements can influence genome organization across different levels. All metrics and contact maps are derived from the predictions of a model trained on HCT116 with only sequence and ATAC-seq as inputs.

As cancerous cell lines exhibit stronger response to perturbation in enhancers compared to normal cells (Supplementary Fig. 16), we next limit our analysis to enhancer regions. **Fig. 4c** compares the averaged curves of the differential insulation scores at 256-kb scale upon silencing enhancers in HIS-subset for HCT116 and its gene-edited variants. An evidently wider peak width of the differential insulation is obtained for HCT116^ΔCTCF^ compared to HCT116 and other variants (**Fig. 4d**), suggesting that the disruption in CTCF boundaries could increase the impact range of local loop formation/destruction. To further check if enhancers within the HIS- subset are of cross-scale impact, we compare enhancers within the HIS-subset obtained at 256-kb and 32-Mb scale in **Fig. 4e**. More than one-third of the enhancers are overlapped at two scales for HCT116 and its variants, confirming pivotal regulatory elements can influence genome organization across different levels. Interestingly, promoters corresponding to these overlapped enhancers with cross-scale impact are more enriched of moderate CpG content (**Fig. 4f**), and more than 90% of these regulatory elements are located at CGI forest regions (**Fig. 4g**). These genes are more likely to be associated with development, mediated by Polycomb interactions^34^. The results are in accord with low level of H3K27me3 marks and moderate expression level of the corresponding genes within HIS-subset (Supplementary Fig. 15). In addition, greater compartment flipping is observed when the corresponding sequence is randomly permuted with segment sizes down to 16 bp (Supplementary Fig. 12) compared to silencing the 1-kb ATAC signals alone, which indicates intrinsic sequence features^23^ (e.g. CpG content) play a key role in maintaining compartment integrity.

### Cancer-induced alterations in genome organization across different scales

Cross-scale correlations could exist for other cancerous cell lines or normal tissues. At loop level, the total number of loops is higher in cancerous cells than in their normal counterparts (**Fig. 5a**). However, higher ratios of loops associated with enhancer-promoter interactions are obtained in normal tissues compared to cancerous cells (**Fig. 5b**). The results indicate aberrant formation of loops enriched with H3K4me1 signals in cancerous cells may disrupt normal enhancer-promoter interactions. We also group detected loops into main-loops and their sub-loops based on their coordinates (Methods). As shown in **Fig. 5c**, larger size of main-loops implies greater loop valencies (i.e. number of sub-loops within the main-loop), but cancerous samples do not always have greater loop valencies compared to their normal counterparts.

**Fig. 5.**
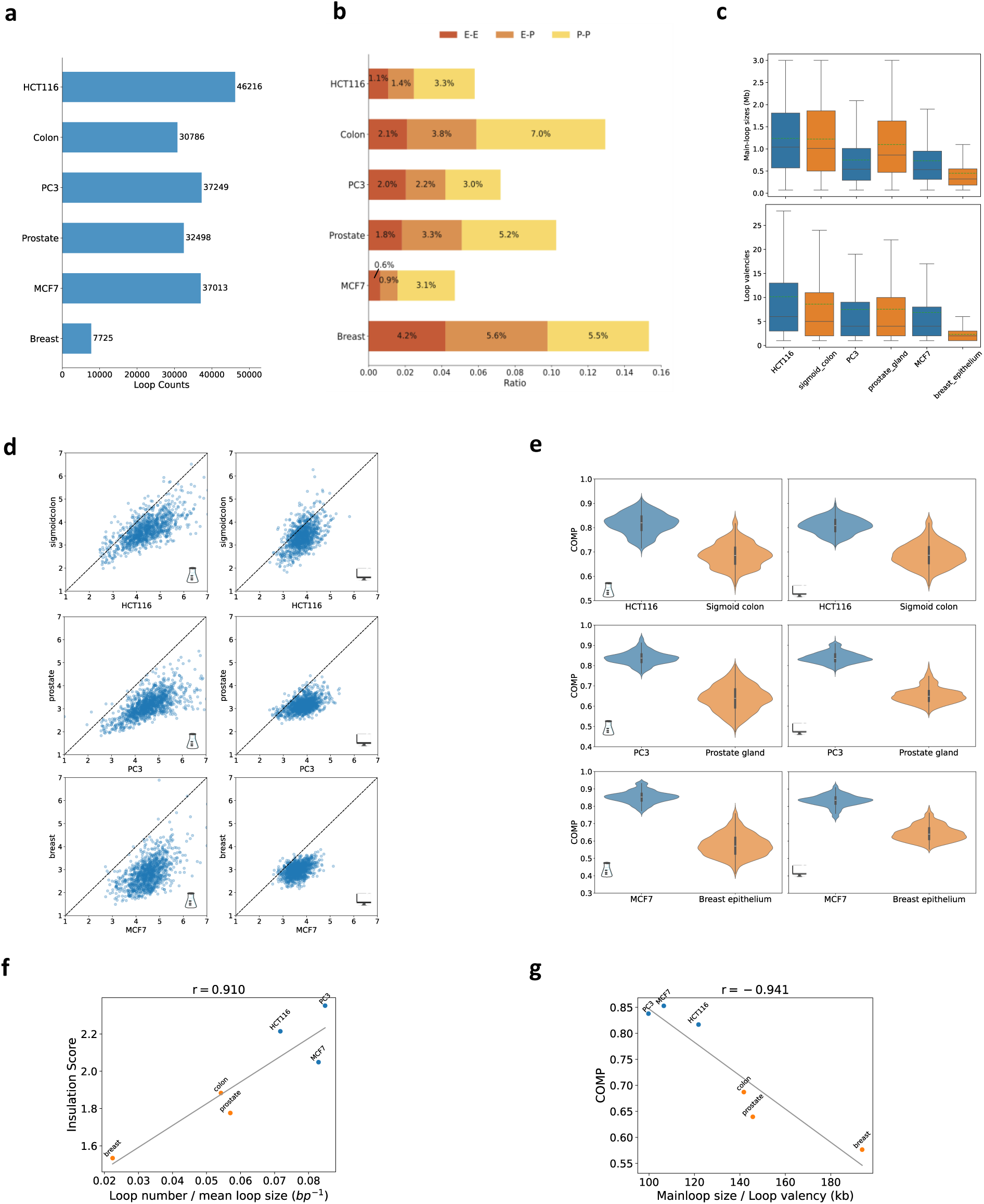
| Comparative analysis of genome organization in normal and cancerous cells across loop, TAD, and compartment levels. a,b, Comparisons of loop counts (**a**) and loop category ratios (**b**) across various cancer and normal samples. Loops are categorized into promoter-promoter (P-P), enhancer-promoter (E-P) and enhancer-enhancer (E-E) based on their coordinates. **c,** Comparisons of main-loop sizes (upper panel) and valencies (lower panel) in cancer cell lines versus normal tissues. **d,** Experimental (left panels) and predicted (right panels) insulation scores at TAD boundaries derived from 8-kb contact maps and for cancerous and normal samples. **e,** Compartmentalization degree (COMP) obtained from experimental (left panels) and predicted (right panels) matrices for normal and cancerous cells derived at 32-Mb scale. **f,g,** Correlations across different scales for cancer and normal samples. Insulation scores correlate positively to loop counts per unit length **(e)**, while COMP correlate negatively to the mean size of loops per unit valency (**f**).

Cancerous cell lines also exhibit higher insulation scores at TAD boundaries compared to their normal counterparts (**Fig. 5d**), suggesting enhanced sub-loop formation and intra-TAD contacts upon carcinogenesis ^35^. Genome-wide compartmentalization degree (COMP) for normal and cancerous cells, evaluated at the 32-Mb scale, are shown in **Fig. 5e**. Both experimental maps and predicted results from HiCGen indicate an increase in COMP after cell carcinogenesis. In fact, the insulation score correlates positively with loop counts per unit length (**Fig. 5f**), with a Pearson coefficients *r* =0.91. The compartmentalization degree correlates negatively with the mean size of loops per unit valency (**Fig. 5g**), with Pearson coefficients *r* =-0.94. These correlations indicate newly formed loops in cancerous cell lines may exert profound influence on chromatin architecture at TAD and even at compartment levels.

### Selective perturbation on regulatory elements related to CRC

To investigate the impact of regulatory elements active in cancerous cell on normal tissue, we conducted *in silico* perturbations on sigmoid colon by activating selected regions corresponding to enhancers in HCT116 (Methods). As shown in **Fig. 6a**, although the compartmentalization degree remained largely unchanged following the perturbation, compartment flipping was observed in regions with emerging enhancer activities. Notably, compartment flipping did take place at the same genome position in HCT116 with respect to sigmoid colon (Supplementary Figs. 17). An enlarged differential map of the flipped region (chr4:136.1-140.3 Mb) at higher resolution revealed alterations in TAD boundaries due to the perturbation (**Fig. 6b**). Specifically, the slight increase in the activity of limited enhancers led to a decrease in insulation strength proximal to SLC7A11 and SETD7, accompanied by enhanced intra-TAD contacts. The enhanced activity of SLC7A11 also exhibits an antagonistic relationship with the PCDH18 locus, resulting in a slight increase in insulation in its vicinity.

**Fig. 6.**
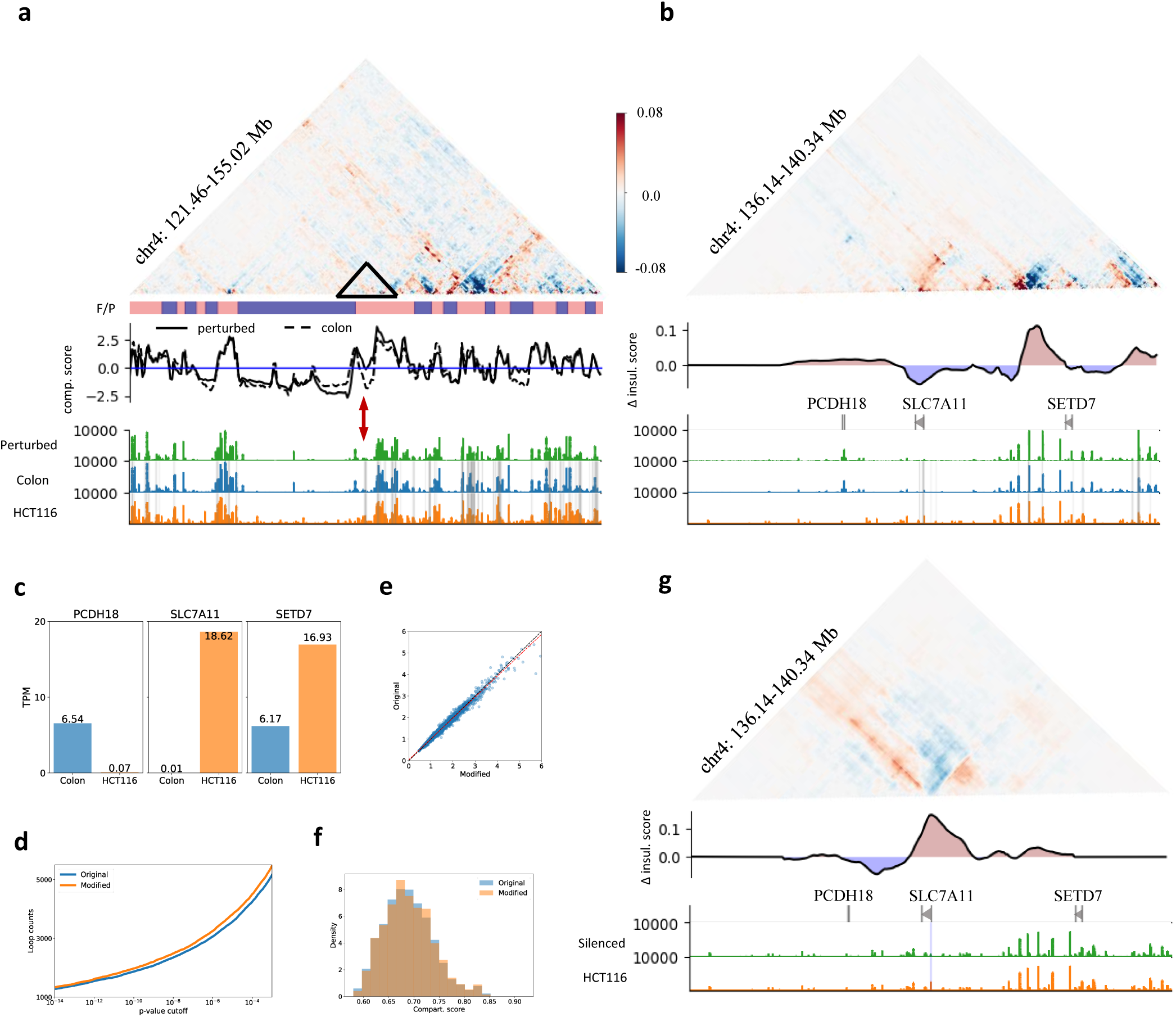
| Selective perturbation on regulatory elements that exert cross-scale impact on genome organizations during carcinogenesis. a,b, Differential contact maps between perturbed and original sigmoid colon for a selected region on chr4, along with CGI forest/prairie domains (F/P), compartment scores at 128-kb resolution (**a**) and differential insulation scores of the corresponding matrices at 16-kb resolution (**b**). The ATAC-seq inputs were ontogenically perturbed by a 1.1-fold increase on selected regions corresponding to enhancers of HCT116. The original and modified ATAC-seq signals are colored in green and blue respectively. Gray vertical lines represent enhancer regions of HCT116. **c,** Comparisons of total RNA-seq TPM for PCDH18, SLC7A11, and SETD7 in sigmoid colon and HCT116. Changes of TPM scores correlate well to the differential contacts and insulations of the corresponding genes upon perturbation. **d-f,** Predicted results of loop counts as a function of p-value cutoff (**d**), insulation scores (**e**) and compartmentalization scores (**f**) between original and perturbed colon genome. **g,** Differential contact maps between silenced and original HCT116 for the same region on chr4, along with the differential insulation scores at 16-kb resolution. Although only one selected ATAC-seq peak corresponding to a 1-kb region centered at the transcription start site (TSS) of SLC7A11 was silenced, it led to an inverse change in contact map and insulation with respect to enhancer activation.

Interestingly, we do observe the upregulation of SLC7A11 and SETD7 and downregulation of PCDH18 in HCT116 when benchmarked against normal sigmoid colon tissue ^33^ (**Fig. 6c**). Both SLC7A11 and SETD7 were reported to play crucial roles in the progression of colorectal cancer (CRC) ^36–41^, while PCDH18 was reported to be frequently inactivated by promoter methylation in CRC ^42^. Despite the pronounced changes in specific TAD boundaries and compartments, the genome-wide insulation score (**Fig. 6e**) and compartmentalization degree (**Fig. 6f**) remained largely unchanged. The modest increase in the number of loops (**Fig. 6d**) induced by the newly activated enhancers was sufficient to perturb the genome into an "oncogenic state", potentially accompanied by subtle alterations in gene expression profiles.

We also selectively perturbed single regulatory element with the corresponding ATAC-seq signals (Methods). **Fig. 6g** presents the effect of silencing the promoter of SLC7A11 in HCT116, which is also one of the pivotal regulatory elements screened out in genome-wide perturbation (**Fig. 4**) with high impact on genome organization. Although only a 1-kb region centered at transcription start site (TSS) of SLC7A11 was silenced, it led to an inverse change in contact map and insulation with respect to enhancer activation in sigmoid colon shown in **Fig. 6b**. We also observed augmented contacts between enhancers of SLC7A11 with neighboring promoters, suggesting the hijacking role of SLC7A11 in HCT116. Supplementary Fig. 11 showcases how perturbations in other genomic regions affect chromosome structure. Activating selected enhancers invariably induces gene-expression-related changes in genome organizations, exemplifying HiCGen as a tool for virtual profiling of pivotal regulatory elements.

## Discussion

The deep learning architecture, HiCGen, is designed to predict genome organizations across diverse cell types and across spatial scales. This model utilizes genome sequence data and genomic features as inputs, enabling *de novo* prediction and perturbation analysis. The output matrices, generated at various resolutions, can be subjected to further downstream tasks, such as 3D genome modeling, comparative and differential analysis. As an illustration, Supplementary Figs. 5-7 respectively compare loops, TADs and compartments detected from experimental and predicted contact maps.

One notable advantage of HiCGen is its scalability, which renders it suitable for interpreting correlations between short-range and long-range interactions of the genome. The model reveals robust correlations across loop, TAD and compartment levels consistent with experimental observations, indicating HiCGen has captured the underlying physical principles governing genomic interactions. The model also accurately predicts coherent changes in both insulation and compartmentalization degree upon editing selected genes in HCT116. According to polymer theory, this cross-scale correlation could stem from the steric repulsion between loop domains ^12^.

For instance, cancerous cell lines typically exhibit a greater number of sub-loops and larger size of main-loops compared to normal cells (**Fig. 5**). These newly emerged sub-loops lead to an increased insulation score and compartmentalization degree in cancerous cell lines relative to normal cells, albeit at the cost of weakened long-range contacts. Our perturbation analysis, as depicted in **Fig. 4**, confirms pivotal role of certain regulatory elements with cross-scale impact on genome organizations. Through either regional or global perturbations on these regulatory elements (Supplementary Fig.11 and **Fig. 6**), we obtain changes in genomic contacts, TAD insulations and even compartment flipping consistent with experimental observations of carcinogenesis-associated changes. We also observed correlations between perturbed structures and expression profiles (**Fig. 6**), which may assist in predicting the latter making use of chromatin structures. We may also perturb ChIP-seq signals of the target sample. For instance, Supplementary Fig.11 showcases enhanced CTCF enrichment at binding sites leads to a decrease in long-range interactions and the collapse of certain fragments with specific orientations of CTCF motifs. The perturbation analysis unravels mechanistic links between epigenetic dysregulation, chromatin reorganization and transcriptional outcomes. Such correlations may provide therapeutic strategies targeting genome misfolding (e.g., correcting oncogenic loops). Additionally, integrating perturbative simulations with multi-omics data could refine synthetic genome design or CRISPR-based editing by forecasting structural consequences.

While HiCGen demonstrates the capability to predict Hi-C maps across cell types at various resolutions, it is not without limitations. Currently, the maximum window size is 32 Mb. To improve efficiency when dealing with even longer sequence lengths, the transformer architectures could be replaced with Mamba ^43^ or Hyena ^44,45^. A lightweight 256-kb model based on HyenaDNA has been trained and compared with HiCGen outputs (Supplementary Fig. 18). Furthermore, structural variants (SV), particularly in cancerous cell lines, such as chromatin breaks, rearrangements, and copy number alteration, have not been adequately characterized^46^.

Training the model with pre-determined SV tags^47,48^ could be a feasible solution to this issue.

## Materials and Methods

### Data preprocessing

The genomic sequences were derived from the GRCh38/hg38 reference genome. We performed five-channel one-hot encoding of the genomic bases ‘ACGT’ and the unknown base ‘N’. The sequencing data for ChIP-seq and ATAC-seq experiments were obtained from the ENCODE data portal (https://www.encodeproject.org/). Details on accession number are listed in Supplementary Table 2-3. Firstly, FASTQ files were subsampled from the merged data of raw replicates. These files were subsequently processed by Seq-N-Slide^49^ protocol to generate the bigWig outputs. For ATAC-seq signals, FASTQ data from different replicates were subsampled down to 40M reads and then processed through Seq-N-Slide ATAC-Route. For ChIP-seq signals, FASTQ data for both experimental and control groups were subsampled down to 30M reads, and then processed through the ChIP-Route. The final outputs are in units of -log10(p-value) for ATAC-seq signals and log2(fold-change to control) for ChIP-seq data. We also take metrics on output files for quality assessment. Tracks with mapped reads ratio less than 80% or duplicates ratio larger than 15% were not taken into account.

The intact Hi-C data were derived from the call sets from the ENCODE portal with identifiers listed in Supplementary Table 2. All data were mapped to hg38 in cool format ^50^. ICE normalization ^51^ with default parameters was performed to the unbalanced data to remove biases. Bins with fewer than 10 nonzero counts were removed from the balanced matrix. Adaptive coarse graining procedures ^52^ were applied to the contact matrices to smooth the low-coverage areas at 1-kb resolution. The average contact frequency as a function of genomic separation over all chromosomes was determined by cooltools^52^.

### Model architecture

The HiCGen is of a typical encoder-decoder framework. The encoder takes the sequence and genomic features as inputs, and increases the represented length from single nucleotides down to 1 kb for each bin by deploying nine blocks of 1D convolutional neural network (1D-CNN). The 1D-CNN module is used mainly for reducing the spatial dimension and increasing channels of each token. The scaling module for the TAD-level model (SwinT-4Mb) is based on the Swin-transformer architecture ^22,23,55^. SwinT-4Mb consists of five consecutive SwinT blocks with a window size 𝑠_𝑤𝑖𝑛_=32. The token length is halved after each SwinT block, and the channel dimension is doubled after the third SwinT block (Supplementary Fig. 1). SwinT-4Mb outputs contact maps at 1-kb, 2-kb, 4-kb, 8-kb and 16-kb resolutions, each with a window size of 256×256 bins. The scaling module for the compartment-level model (SwinT-32Mb) is built upon the 4-Mb model, with four additional SwinT layers (Supplementary Fig. 1). SwinT-32Mb outputs contact maps at 16-kb, 32-kb, 64-kb, and 128-kb resolutions. The decoder part is kept as a 2D convolutional neural network (2D-CNN). At largest predicting window (i.e. 16-kb resolution in SwinT-4Mb and 128-kb resolution in SwinT-32Mb), the 2D-CNN module contains five dilated residual blocks. For decoders at finer resolutions, prediction from the coarser resolution corresponding to the same region is up-sampled and incorporated into the input to the 2D-CNN module.

We also implement a hybrid model of 256-kb genomic length based on Hyena and SwinT architectures (Hyena- SwinT-256kb) for comparison (Supplementary Fig. 18). We only implement Hyena module for the first two layers of Hyena-SwinT-256kb, with all default setting inherited from HyenaDNA ^44^.

### Training

The model was trained on a GPU cluster with quad NVIDIA Tesla A100 80GB GPUs. The data-loader generates training data, including sequence, genomic features and Hi-C data at 1-kb resolution for SwinT-4Mb and 16-kb resolution for SwinT-32Mb. For prediction tasks, DNA sequence, ATAC-seq and CTCF ChIP-seq signals were implemented as model inputs. For perturbation tasks (Fig. 4), Only DNA sequence and ATAC-seq signals were implemented as model inputs. The data were obtained by uniformly sampling the genome from training chromosomes. Chromosome were split into training, validation and test set based on the Chromosome ID. The entire chr10 was assigned to the validation set while the entire chr15 was assigned to the test set. The rest autosomes were assigned to the training set. We further obtained contact matrices at lower resolutions by average pooling the original Hi-C data in 2D. Contact matrices were down-sampled to meet the prediction windows at multiple resolutions for the 4-Mb and 32-Mb models. To reduce overfitting, we replaced about half of the randomly selected forward- sequences with their reverse-complement counterparts during the training process. Gaussian noise with a mean of zero and a standard deviation of 0.1 were added to input signals.

The model was trained to predict the log-fold contact maps over genomic-distance-based background. We trained the model with Adam optimizer using a cosine annealing schedule, with an initial learning rate of 0.001 and batch size of 4. For both SwinT-4Mb and SwinT-32Mb models, the output matrices were compared with the experimental Hi-C map via a mean squared error (MSE) loss. Checkpointing was used for lower memory consumption. Particularly, the SwinT-32Mb model was trained in two stages: first, the model was trained up to 4 Mb, similar to that for the SwinT-4Mb model; then, the encoder and the first five consecutive Swin-transformer blocks were frozen and used in the later-stage training of the following four SwinT blocks and decoders.

To enhance prediction accuracy, non-readable regions (including telomeres or centromeres) were masked. The model was trained on DNA sequence, CTCF ChIP-seq and ATAC-seq profiles of the GM12878 cell line. We also provided models trained on sigmoid colon and PANC-1 for better performance in predicting contact maps of normal tissues and cancerous cell lines respectively.

### Evaluation

For evaluating the model trained on GM12878, multiscale predictions of the genome interaction matrices were compared with the experimental data for the test chromosomes. For *de novo* prediction of contact matrix for other cell lines, we replaced CTCF ChIP-seq and ATAC-seq with the corresponding signals from the specific cell line while keeping the same reference DNA sequence.

Pearson correlation between the predicted and experimental contact matrices was computed. We also compared background-subtracted correlation and distance-stratified correlation, both of which were calculated as Pearson coefficient between the shifted diagonal line of prediction and ground truth for evaluating long-range interactions.

We presented comparison with previously released Hi-C predictors such as DeepC ^22^, Orca ^23^ and C.Origami ^24^ (Supplementary Fig. 2). Since DeepC was trained to predict a 1-Mb window of contacts, we restricted regions for comparison to 1-Mb blocks. We selected 1-Mb regions in a sliding window with 0.5-Mb overlap between neighboring regions, which were shared to each model for prediction. For better relevancy for benchmarking tests, we limited our comparisons to unseen (i.e. test set) chromosome of the target cell line in training. For DeepC, chr16 and chr17 were used with the model trained with GM12878; for Orca, chr9 and chr10 were used with the model trained with HFF; for C. Origami, chr15 was used with the model trained with IMR90; for HiCGen, chr15 was used with the model trained with GM12878. We performed additional genomic-distance-based normalizations to the predicted results (and also the target maps) of C.Origami to match the training targets of DeepC, Orca and HiCGen. To align with the 8-kb resolution of the prediction of C.Origami, we selected the 2-Mb window in Orca and HiCGen and cropped out the centering 1-Mb window for comparison. Since DeepC was trained to predict rotated contact maps at 5-kb resolution, we performed cropping, rescaling and 45 degree rotation to its outputs.

We also presented cross-scale comparisons between HiCGen and Orca. Since both DeepC and Orca do not allow cross-cell-type predictions, benchmarking performance on *de novo* prediction of unseen cell types was compared only between HiCGen and C.Origami (Supplementary Fig. 3-4).

### Identification and evaluation of loops

Loop calling for normalized Hi-C map scaled to 10-kb-resolution was conducted using Peakachu^53^, which employs a Random Forest classification framework to identify chromatin loops. Detected loops are further categorized into promoter-promoter (P-P), enhancer-promoter (E-P), enhancer-enhancer (E-E) and other type based on the coordinates of the two loop boundaries.

Loops are grouped into main-loops and their sub-loops based on their coordinates^12^. Loop *α* is considered to be a sub-loop of loop *β* only if the coordinates follow:

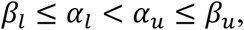

where 𝛼_𝑙_, 𝛼_𝑢_, 𝛽_𝑙_, 𝛽_𝑢_ respectively denote the lower- and upper-bound coordinates for loop *α* and loop *β.* The loop valency for a given main-loop is defined as the number of sub-loops within the main-loop. To evaluate the model performance in predicting loop domains, we computed the area under the receiver operating characteristic curve (AUROC) and overlap between loops called from experimental Hi-C and loops called from predicted Hi-C (Supplementary Fig. 5). We also tested loop callers such as FitHiC^54^ and Mustache^55^ at various resolutions for comparison, which gave more false positive calls below 2-kb resolution than those by Peakachu^53^.

### Identification and evaluation of TADs

TAD boundaries for the normalized Hi-C contact map were identified by FAN-C^56^. The Hi-C resolution was set to 25 kb with a window size of 1 Mb. The calculation of delta vector takes into account 3 bins upstream and 3 bins downstream for each locus and call minima of insulation scores as TAD boundaries.

The degree of insulation (INSUL) were characterized by the insulation score ^57^ at TAD boundaries. Within a window of 50 bins upstream and 50 bins downstream, the insulation score for given point is calculated as the ratio of left (L) and right (R) region average intensity and the middle (M) region average intensity, which can be written as:

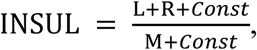

where the constant equals to the average intensity of the Hi-C contact map at given resolution.

### Identification and evaluation of compartments

Hi-C contact matrix was regarded as an adjacency matrix, and spectral clustering and Linear Discriminant Analysis dimensionality reduction^58^ were performed on iced Hi-C at 32-kb, 64-kb and 128-kb resolutions to identify compartments^59^. Eigenvalue decomposition was performed over Laplacian matrix, which was computed corresponding to the Hi-C matrix of each individual chromosome. The top 50 eigenvectors were chosen for linear discriminant analysis (LDA) dimensionality reduction. The dimensionally reduced output serves as indicator vectors for compartments A and B, where a positive or negative sign indicates compartment A or compartment B, respectively.

The dimensionless order parameter, COMP, is introduced to measuring the compartmentalization degree of the 32 Mb genome (with log-fold contact maps over genomic-distance-based background), which can be written as:

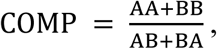

where AA, AB, BA, BB respectively denote contact probabilities in the corresponding quadrant. We further rescale COMP by:

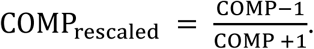

Now COMP_rescaled_ = 1 denotes contacts are only within compartments but none across, whereas COMP_rescaled_ = −1 denotes contacts are only across compartments.

### Two-dimensional layout of chromatin structures

To visualize the chromatin structures, we first performed CTG^60^ (Hi-C To Geometry) on iced Hi-C contact matrix to eliminating systematic bias and get reliable spatial distance matrix. Then we used the Fruchterman-Reingold algorithm and Monte-Carlo sampling based on CTG matrices to generate reproducible two-dimensional force-directed layouts ^59^.

### Categorizing genome fragments

We divided the genome into fragments of 1-kb length for perturbation tests. The 1-kb fragments can be categorized according to their genomic function, namely gene bodies, promoters, enhancers and other regions. The promoter region was defined as a window spanning 5 kb upstream and 500 bp downstream of the gene transcription start site. Enhancers were derived from ENCODE annotations, which were defined by the activity-by-contact method^61^ utilizing DNase-seq and ChIP-seq signals of the corresponding cell line.

On compartment scale, the genome can be categorized into CGI forest regions and prairie regions, which, respectively, correspond to chromosomal segments with either rich or poor densities of CpG islands^62^. The relative enrichment of regulatory elements of cross-scale impact (**Fig. 4e-g**) in CGI forest/prairie is defined as the ratio of the element count to the total number of genes within CGI forest/prairie regions. Genes can also be categorized based on their CpG density proximal to TSS^34^. Correspondingly, the relative enrichment of selected genes with different CpG density is defined as the ratio of the count of selected genes to the total number of genes within low, moderate or high CpG content.

### Regional perturbation

We perturbed regional activity of the genome with the corresponding ATAC-seq signals. Regional perturbation of the genome can be applied to any given or selected genomic regions with a model trained without ChIP-seq inputs. To screen out regions with high impact on genome organizations, we first delimited the genome into fragments of 1-kb length. A chromosome-wide comparison of the impact of silencing 1-kb ATAC-seq signals on contact matrices was performed. The impact score in the screening experiment is defined as the mean absolute difference between predictions prior to and subsequent to the silencing of the signals^24^. Note that the impact scores could vary significantly across scales.

At loop/TAD scale, silencing 1 kb of ATAC-seq signals leads to either the decreased or increased of insulation. While a decrease in insulation scores can be attributed to the loss of protein-mediated or enhancer-promoter interactions, an increase in insulation scores may result from the compartment flipping of the silenced region.

The flipping can establish an insulating boundary and disrupt enhancer-promoter interactions at both ends (blue arrows in Fig. 4a). For each type of silencing tests, we selectively analyze differential insulation curves for the top 10% fragments with highest impact score (HIS-subset) at 1-kb and 4-kb resolutions. Specifically, the differential insulation curves within HIS-subset are categorized into positive and negative ones based on the sign of peak values. The peak value is defined as either the maximum or minimum of the insulation curve. The impact width of the perturbation on insulation of the fragment is defined as the maximum bin number in proximity to the peak position in both directions without changing the sign.

The regional activation of carcinogenesis-associated enhancers in sigmoid colon was modeled by directly replacing ATAC-seq signals of sigmoid colon with ATAC-seq signals of HCT116 enhancers within selected 1- Mb, 256-kb or 1-kb fragment (Supplementary Fig. 11). The original ATAC-seq peaks were replaced only when the replaced peak was greater than the original one. The regional deactivation of carcinogenesis-associated regulatory elements in HCT116 was performed by silencing chosen region of ATAC-seq signals corresponding to either enhancer or promoter of HCT116.

Regional perturbations on CTCF binding enrichment were trained with model without ATAC-seq inputs. The activation of insulation boundaries was modeled by adding CTCF ChIP–seq peaks in proximity to the CTCF motifs within the modified segment. Each added peak is of Gaussian distribution, with peak position right at the middle of the CTCF motif. The peak value equals to 1000, and the peak width equals to the length of the CTCF motif extended by 30 bp in both directions. The original CTCF ChIP–seq signals are replaced only when the enhanced value is greater than the original one. If there is no inactivated CTCF motifs within the selected region, we create CTCF boundaries through inserting CTCF motifs at specific position of the segment^63^.

We also test the effect of genome sequence on perturbation with random permutation. The sequence within the perturbed region was first divided into segments of 16 bp. Then, the order of the sequence segments was randomly permuted to disrupt their compartmental identities.

### Global perturbation

For evaluating the effect of average CTCF binding affinity or chromosomal accessibility on compartmentalization degree, we performed global perturbation of either CTCF ChIP–seq or ATAC-seq signals. To test the effect of CTCF binding affinity, all CTCF ChIP–seq signals were multiplied by a fold-change factor, while ATAC-seq signals remained unchanged. To test the effect of chromosomal accessibility, ATAC-seq signals were multiplied by a fold-change factor, while CTCF ChIP–seq signals remained unchanged (Supplementary Fig. 12) .

To evaluate the effect of activating carcinogenesis-associated regulatory elements on genome interactions of normal cells, we performed global perturbation of ATAC-seq profiles in enhancers regions of sigmoid colon.

The global activation of cancerous enhancers on sigmoid colon genome was modeled by multiplying the value of ATAC-seq signals at enhancers of HCT116 derived from ENCODE with a 1.1-foldchange factor (**Fig. 6**).

## Acknowledgements

This work was supported by National Science and Technology Major Project (No. 2022ZD0115001), National Natural Science Foundation of China (92353304, T2495221) and New Cornerstone Science Foundation (NCI202305). Computations were carried out at Changping Laboratory Supercomputing Center.

